# Using Culture-Enriched Phenotypic Metagenomics for Targeted High-Throughput Monitoring of Clinically-Important Fraction of Beta-Lactam Resistome

**DOI:** 10.1101/2022.04.06.487307

**Authors:** Zhiguo Zhang, Guoqing Zhang, Feng Ju

**Affiliations:** Key Laboratory of Coastal Environment and Resources of Zhejiang Province, School of Engineering, Westlake University, 18 Shilongshan Road, Hangzhou 310024, Zhejiang Province, China; Institute of Advanced Technology, Westlake Institute for Advanced Study, 18 Shilongshan Road, Hangzhou 310024, Zhejiang Province, China

**Author notes:** **Corresponding to:** Dr. Feng Ju (Assistant Professor), Address: Environmental Microbiome and Biotechnology Laboratory (EMBLab), Westlake University, 18 Shilongshan Road, Hangzhou 310024, China, (lab), 571-87380995 (office).

**Keywords:** Phenotypic Metagenomics, Carbapenemases, Extended-Spectrum Beta-Lactamases (ESBLs), Wastewater Treatment Plants, Microbiome

## Abstract

The high bacterial community complexity greatly hinders the monitoring of clinically prevalent extended-spectrum beta-lactam resistant bacteria, which are usually present as rare but key populations involving in the environmental dissemination of clinical resistance. Here, we introduce culture-enriched phenotypic metagenomics that integrates culture enrichment, phenotypic screening and metagenomic analyses as an emerging methodology to profile beta-lactam resistome in a municipal wastewater treatment plant (WWTP) and its receiving river. The results showed that clinically prevalent carbapenemase genes (e.g., NDM and KPC family) and extended-spectrum beta-lactamase genes (e.g., CTX-M, TEM and OXA family) considerably existed in the WWTP and showed prominent potential in horizontal dissemination. Strikingly, carbapenem and polymyxin resistance genes co-occurred in highly virulent *Enterobacter kobei* (MCR-1 and NDM-6) and *Citrobacter freundii* (ArnA and NDM-16) genomes. Overall, this study exemplifies phenotypic metagenomics for high-throughput surveillance of targeted fraction of clinically-important resistome and substantially expands current knowledge on extended-spectrum beta-lactam resistance in WWTPs.

## Introduction

Extended-spectrum cephalosporins and carbapenems are of critical importance to the treatment of nosocomial infections caused by multi-antibiotic resistant human bacterial pathogens (HBPs) ^1^. However, the clinical outcomes of these key antibiotics are seriously compromised by the spread of extended-spectrum beta-lactamases (ESBLs) and carbapenem hydrolyzing enzymes called carbapenemases ^2,3^ in clinical HBPs, such as *Pseudomonas aeruginosa, Acinetobacter baumannii*, and *Enterobacteriaceae*, resulting in numerous outbreaks of bacterial infection and alarming patient mortality throughout the world in recent decades ^3-5^. Municipal wastewater treatment plants (WWTPs) continuously receive wastewater including human excrements and hospital wastewater discharges, which potentially carry a diverse amount of antibiotic resistance genes (ARGs) and clinical HBPs ^6-9^. However, when introduced into an environmental microbiome, ARG-carrying HBPs and other enteric bacteria are often metabolically inactive, less competitive, and passively diluted to ultra-low abundance that can easily fall below detection limits of direct metagenomic sequencing. Therefore, the fate and resistome profiles (i.e., the collection of all ARGs ^10^) of clinically-important but mostly rare carbapenem and cephalosporin resistant bacteria, especially for HBPs, in WWTPs and the receiving rivers still remain elusive.

For decades, culture-dependent methods that combined bacterial colony isolation with conventional PCR have been commonly used for the investigation of specific antibiotic resistance phenotypes in targeted bacterial populations ^11-13^. However, such classic methods are not only extremely limited in their detection throughput, but also high-cost and labor-intensive (e.g., tedious experimentation), especially inefficient for large-scale surveillance. More recently, culture-independent high-throughput qPCR ^14,15^ and metagenomic sequencing ^6,9^ are increasingly utilized to profile ARGs, greatly contributing the surveillance of environmental resistomes. However, high-throughput qPCR, which relies on primers selection and amplification as conventional PCR does, captures only part of environmental resistomes and fails to fetch other key traits of ARGs, such as gene structure, host information and functional activities. Metagenomic sequencing well overcomes some of the above limitations and realizes high-throughput profiling of ARGs, while, due to extremely complex genetic background, it is difficult to fetch the desired information from complex environmental samples in an affordable way, as ultra-high sequencing depth is usually required for effective recovery of genomic contents of rare species, such as carbapenem and cephalosporin resistant *Enterobacteriaceae* in environmental reservoirs. Collectively, it remains challenging to get a comprehensive understanding on the environmental prevalence, spread or health risks of clinically-important ARGs (e.g., carbapenem and cephalosporin resistance genes) and key features of their hosts based on solely culture-dependent colonies isolation or culture-independent genotyping methods. Therefore, a new strategy and related approaches are urgently needed to address these long-standing challenges.

Taking environmental surveillance of carbapenem and cephalosporin resistant bacteria as an example, we propose and demonstrate an integrated methodology based on culture enrichment, phenotypic screening and metagenomic analysis of environmental microbiomes, namely culture-enriched phenotypic metagenomics (Fig. 1), to explore the fate, genotypic profiles, pathogenicity and taxonomic identity of carbapenem and cephalosporin resistant microbiomes with targeted resistance phenotypes throughout a local WWTP and its receiving river. The endeavors that couple phenotypic and genotypic analyses of antibiotic resistance would offer new insights into the targeted clinically-important fraction (e.g., extended-spectrum beta-lactam resistance) of wastewater beta-lactam resistomes by overcoming the evident limitations of each analysis when applied alone ^6,12,15,16^. Moreover, we also check and discuss the feasibility of culture-enriched phenotypic metagenomics as an alternative basis for developing standardized surveillance methods of environmental resistomes.

**Fig. 1.**
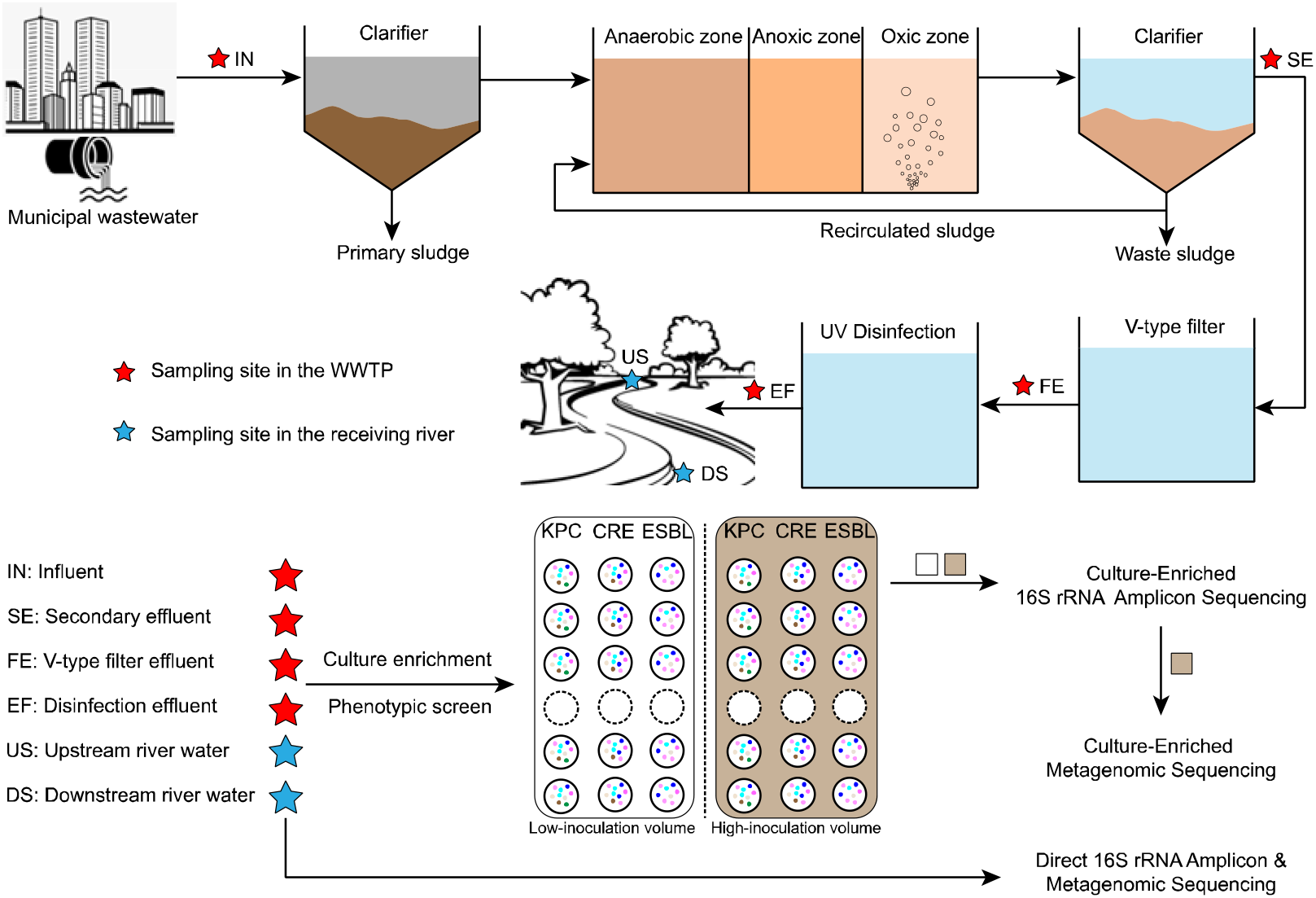
The workflow diagram of culture-enriched phenotypic metagenomics and the tertiary municipal wastewater treatment plant (WWTP) under study. Upper panel: six sampling sites in the WWTP and its receiving river. Lower panel: culture-enriched phenotypic metagenomics that intergrate phenotypic screening, 16S rRNA gene amplicon sequencing, and metagenomic sequencing to analyze the targeted fraction of environmental microbiomes. In brief, the biomass-carrying membranes were incubated in triplicate for each sample on three different selective agars (CHROMagar™, France), that is, the KPC agar for detection of Gram-negative bacteria with a reduced susceptibility to most of carbapenem antibiotics, mSuperCARBA (CRE) agar for detection of Carbapenem-resistant Enterobacteriaceae, and ESBL agar for detection of gram-negative bacteria producing extended-spectrum beta-lactamases, which are widely applied for phenotypic screening of clinical carbapenem and cephalosporin resistance.

## Results

### Bacteriome diversity and WWTP fate of carbapenem and cephalosporin resistant microbiomes cultured from wastewater and river water

The KPC, CRE and ESBL agar media, which have been commonly applied to identify carbapenem and cephalosporin resistant clinical pathogens, were used to phenotypically screen and independently select microbiomes inoculated namely from four wastewater samples (i.e., IN, SE, FE and EF) collected along a tertiary WWTP and two water samples (i.e., US and DS) of its receiving river (Fig. 1). The phenotypically screened culture-enriched samples generated were analyzed based on 16S rRNA gene amplicon sequence variants (ASVs) to taxonomically explore the diversity of carbapenem and cephalosporin resistant bacteria in the WWTP and its receiving river, and further evaluate the influence of inoculation volume so as to optimize biomass yield for culture-enriched metagenomic sequencing. The results showed that the culture-enriched bacteriome composition was greatly differed and distinct among both habitat types (ANOSIM test, R=0.69, *P* < 0.01) and sampling sites (ANOSIM test, R=0.12, *P* < 0.05), (Fig. 2A and 2B). The culture-enriched carbapenem-resistant microbiomes on the KPC agar was dominated by *Pseudomonas* (42.1%, on average hereinafter), *Acinetobacter* (17.7%) and *Stenotrophomonas* (11.8%), and those on the CRE agar by *Escherichia* (10.8%) and *Pseudomonas* (64.6%), while cephalosporin resistant microbiomes selectively enriched on the ESBL agar mainly consisted of *Escherichia* (34.5%), *Klebsiella* (10.5%) and *Pseudomonas* (21.1%). The distribution of extended-spectrum beta-lactam resistant *Pseudomonas* in aquatic ecosystem has been widely described ^17^. The CRE agar was primarily designed to identify carbapenem-resistant *Enterobacteriaceae*, so *Enterobacteriaceae* species (e.g., *Escherichia* and *Klebsiella*) were unsurprisingly more abundant than that on the KPC agar (15.9% vs. 3.8%), (Fig. S1). The enrichment of carbapenem resistant *Enterobacteriaceae* species on CRE agar would facilitate the exploration of their genomic features based on further metagenomic analyses. To check the influence of inoculation volume, the agar plates was seeded with 0.1 to 500 ml of wastewater or river water samples depending on expected biomass through filter membranes (X-axis labels, Fig. 2A). The results showed no significant difference in the composition of culture-enriched bacteriomes between two contrasting inoculation volumes examined (i.e., 0.1 vs. 1 ml for IN, 10 vs. 100 ml for SE, US and DS, and 100 vs. 500 ml for FE; Wilcoxon signed-rank test, *P* > 0.10). Thus, culture-enriched samples with higher culturable biomass (usually with high inoculation volumes) were selected for metagenomic sequencing, so as to better impair the sequence contamination of initial microbial inocula in culture-enriched metagenomic data.

**Fig. 2.**
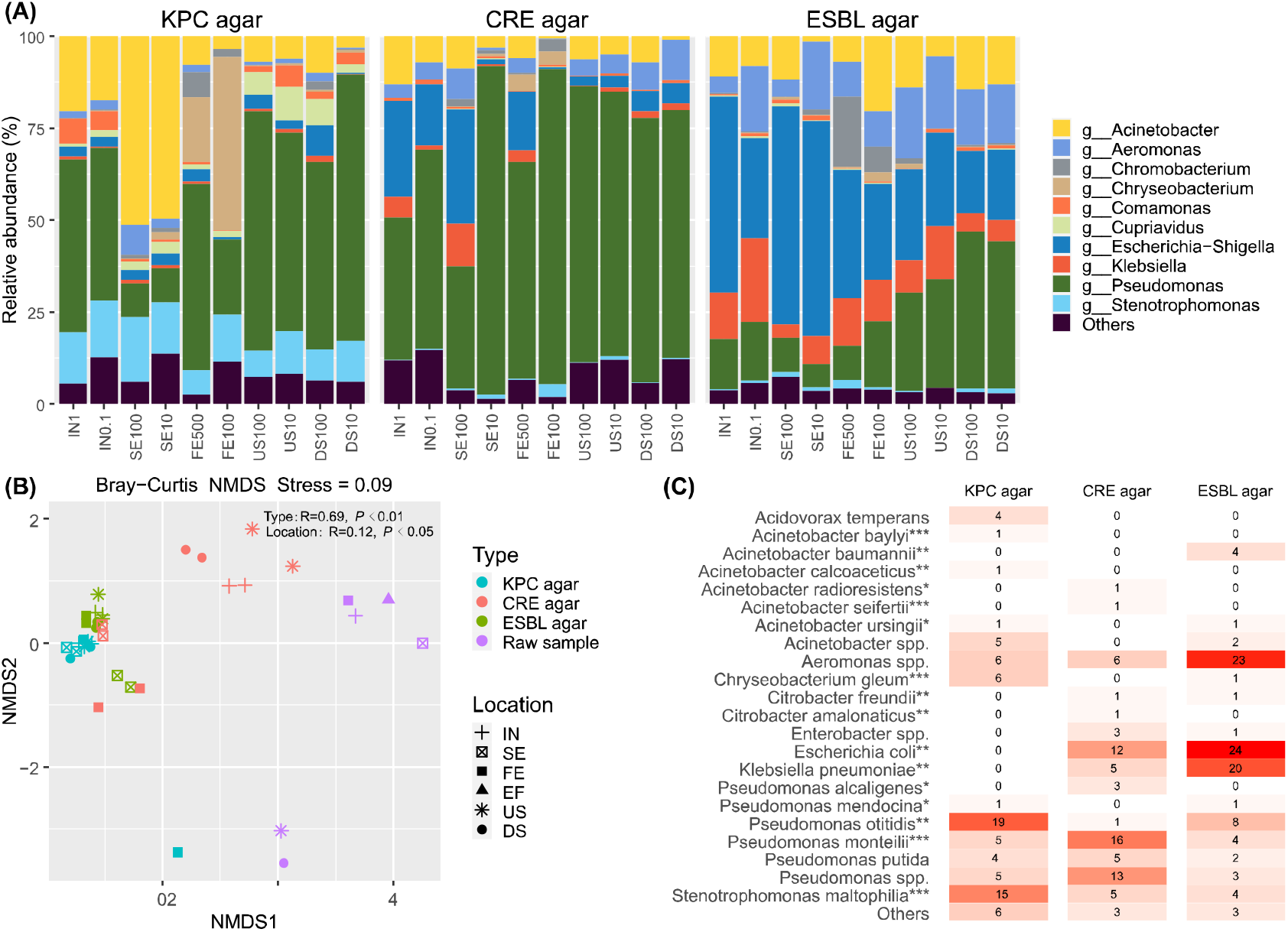
Bacterial community composition and beta-diversity of the culture-enriched and before-culture environmental samples. (A), 16S rRNA gene amplicon sequence variants (ASVs) analsyis of culture-enriched microbiome composition. IN: influent, SE: secondary effluent, FE: V-type filter effluent, US: upstream, DS: downstream, (B), Beta-diversity analysis with non-metric dimentionsal scaling (NMDS). (C), MALDI-biotyping of resistant colonies. Number in the cell denotes no. of isolates, and asterisk denotes clinical importance of emerging global opportunistic human pathogen (***), opportunistic human pathogen (**), and potential human-infectious bacteria (*).

To validate the ability of culture-enriched phenotypic metagenomics on detection of clinically-important but rarely-present extended-spectrum beta-lactam resistant HBPs, MALDI-TOP biotyping was further used to taxonomically assign resistant colonies randomly picked from the selective agar plates (Fig. 2C). In line with the 16S data, the MALDI-TOP biotyping results showed different bacteriome composition between phenotypically screened carbapenem and cephalosporin resistant microbiome originating from wastewater and river water. On the KPC agar and CRE agar, *Pseudomonas* and *Stenotrophomonas* were the dominant species, while *Escherichia* and *Klebsiella* were the predominant ones on ESBL agar. In the 79 KPC-agar bacterial isolates, 29 and 20 isolates were alarmingly assigned to emerging global opportunistic HBPs (e.g., *Chryseobacterium gleum* and *Stenotrophomonas maltophilia*) and opportunistic HBPs (e.g., *Pseudomonas otitidis*), respectively. In contrast, the 102 cephalosporin resistant isolates were most represented by *Aeromonas spp*. (23 isolates) and three opportunistic HBPs, i.e., *Escherichia coli* (24 isolates), *Klebsiella pneumoniae* (20 isolates), and *Acinetobacter baumannii* (4 isolates).

### Resistome diversity and WWTP fate of carbapenem and cephalosporin resistant microbiomes

To link antibiotic resistance phenotype to its associated genotypes in a complex setting of environmental microbiomes, antibiotic resistomes of wastewater and river water samples before and after phenotypic screening were profiled based on read-level estimation of average <u>G</u>ene copies <u>P</u>er bacterial <u>C</u>ell (GPC) ^16,18^. In total, 17 ARG types were identified from six wastewater or river water samples and 15 culture-enriched samples (Fig. 3A). Culture-enriched metagenomic sequencing, even with almost half lower sequencing depth (6.1 Gbp vs. 10.6 Gbp per sample), detected greater diversity and number of ARGs compared with direct metagenomic sequencing (Fig. S2) owing to the designed phenotypic selection. To quantitatively measure the enrichment of ARGs on the selective agar, we calculated selective enrichment factor (SEF) as the metagenomic abundance ratio of a given ARG from after to before phenotypic selection. Despite that KPC, CRE and ESBL agar media were all designed to screen for beta-lactam resistance, resistance types (e.g., multidrug, aminoglycoside, tetracycline, MLS and sulfonamide) other than beta-lactamase genes were also markedly enriched (i.e., SEF > 5) by at least one of the three agar media, as shown by black circles (Fig. 3A), revealing the potential co-selection of beta-lactamase genes with other ARGs in the culture-enriched carbapenem and cephalosporin resistant microbiomes. The co-occurrence patterns between ARGs were further explored by gene arrangement analysis of metagenome-assembled contigs, identifying 443 of 2521 ARG-carrying resistance contigs encoding at least two co-occurring ARGs. For example, one resistance plasmid contig (CFE500_3208), encoding Intl1 integrase, was identified to contain 6 ARGs conferring broad resistance to five classes of antibiotics, namely sulfonamide, rifamycin, chloramphenicol, beta-lactam, and aminoglycoside, which results in spreads of multi-antibiotic resistance once horizontally mobilized (Fig. S3).

**Fig. 3.**
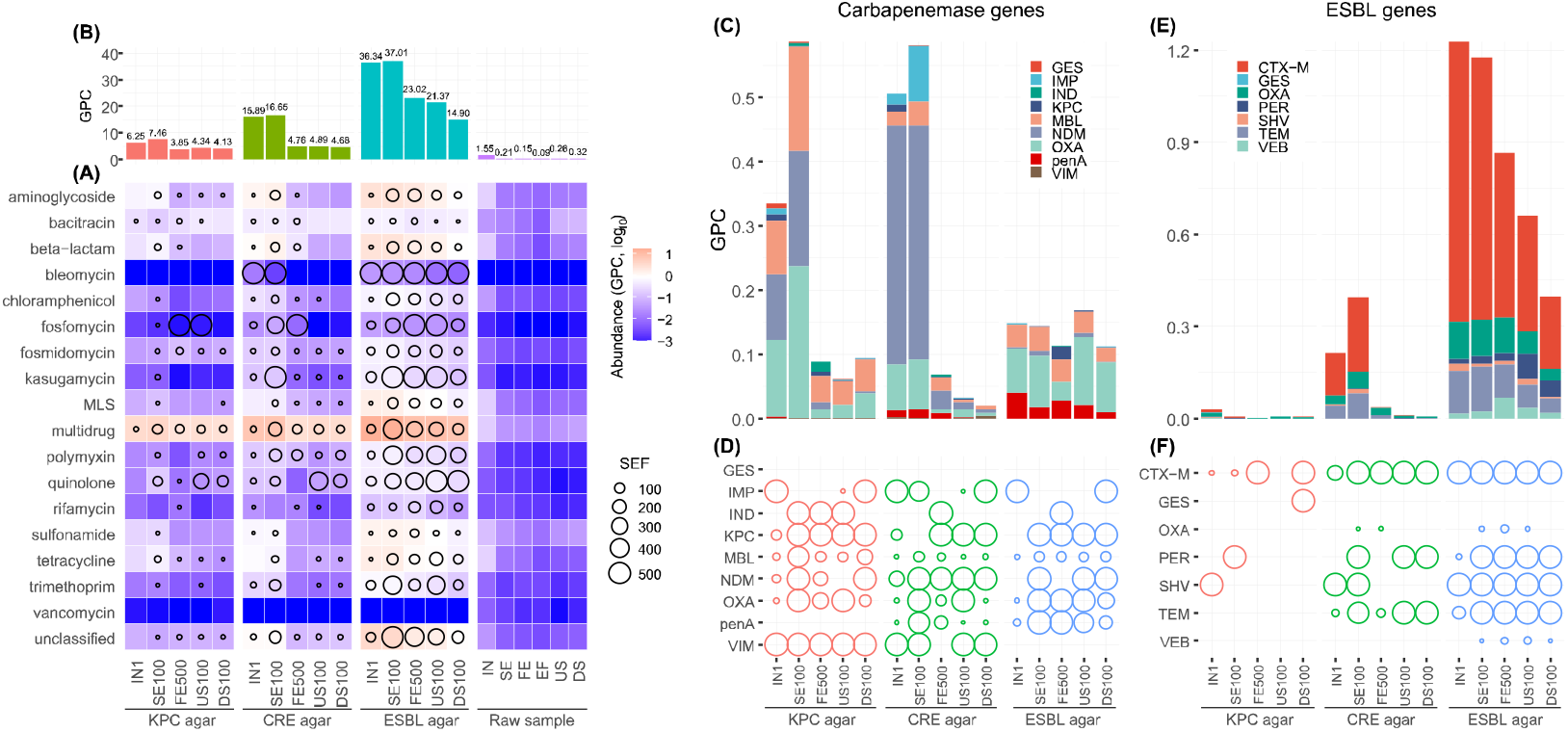
Resistome composition of culture-enriched and before-culture environmental samples. (A), The relative abundance of different ARG types. SEF represents selective enrichment factor of ARGs after relative to before microbiota selection on the agar, which is shown by circles. Circles are shown only when SFE > 5. GPC: gene copies per cell, MLS: macrolide-lincosamide-streptogramin. (B), The total relative abundance of ARGs in each sample. (C) and (E), The composition of beta-lactamase genes on the KPC, CRE and ESBL agars originating from wastewater and river weater bacteria. (D) and (F), The selective-enriched fold of beta-lactamase genes on the KPC, CRE and ESBL agars compared with original water.

We further questioned which subtypes of beta-lactamase genes are responsible for the observed resistance phenotypes on the presumptive carbapenemases-selective (i.e., KPC and CRE) and ESBLs-selective agars. Overall, carbapenemase genes (Fig. 3C) and ESBL genes (Fig. 3E) were dominant in carbapenemases-selective and ESBLs-selective agar, respectively, in line with their acknowledged resistance functions of extended-spectrum beta-lactams (e.g., carbapenems and cephalosporins). On one hand, nine carbapenemase families belonging to class A (i.e., GES, KPC and penA), class B (i.e., IMP, IND, MBL, NDM and VIM), and class D (i.e., OXA) beta-lactamases were detected in carbapenem-resistant microbiomes (Fig. 3C). MBL, NDM and OXA family carbapenemases were the major responsible genes for resistance on KPC agar (0.08, 0.10 and 0.12 GPC in IN, respectively), while NDM genes were solely predominant on CRE agar (0.37 GPC in IN). On the other hand, seven ESBL families belonging to class A (i.e., CTX-M, GES, PER, SHV, TEM and VEB) and class D (i.e., OXA) beta-lactamases were detected in ESBL-agar microbiomes (Fig. 3E), in which the CTX-M family (0.90 GPC in the influent) was the most dominant, followed by TEM family (0.14 GPC) and OXA family (0.12 GPC). Noteworthy, we identified a set of markedly selected beta-lactamase genes (SEF > 5), such as NDM, MBL, KPC and OXA-family carbapanemases on KPC agar and CRE agar (Fig. 3D), and CTX-M, TEM and OXA-family ESBLs on ESBL agar (Fig. 3F). Hypothetically, environmental bacteria harboring these potent ARGs of extended-spectrum beta-lactams (i.e., cephalosporins and carbapenems) should well proliferate on the selective agars. Intriguingly, the dominant beta-lactamase gene families, i.e., OXA, TEM, SHV, CTX-M and NDM, were also the main beta-lactamase types commonly found in clinical pathogens (Table S4), implying the prevalence of clinically relevant extended-spectrum beta-lactamase resistance in wastewater and river water examined.

In raw wastewater, the total abundance of ARGs greatly decreased from 1.55 GPC in the influent (IN) to 0.21 GPC in secondary effluent (SE) and further to 0.09 GPC in the final post-treatment effluent (EF), (Fig. 3B). In contrast, the resistome abundance of culture-enriched microbiomes on three selective agars (6.25, 15.89 and 36.34 GPC on KPC, CRE and ESBL agar, respectively, in IN) showed no decrease in SE (7.46, 16.65 and 37.01 GPC on KPC, CRE and ESBL agar, respectively), and was reduced till after tertiary filtration, by 38.4%, and 70.0% and 36.7%, respectively, to levels comparable to that of the receiving river water (Fig. 3B). Thus, it is not suitable to directly discharge secondary effluent into the environment, as also evidenced earlier for WWTPs in Swiss ^16^ and other European countries ^15^, and tertiary treatment with sand filter plays a critical role on ARGs mitigation of cultivable carbapenem and cephalosporin resistant microbiomes. In line with total abundance of ARGs, carbapenemase genes (Fig. 3C) and ESBL genes (Fig. 3E) in SE were still as abundant as those in IN, and were greatly reduced by tertiary filtration by 73.4 % and 86.6% for KPC and CRE agar-enriched carbapenemase genes, respectively, and 29.6% for ESBL agar-enriched ESBL genes (Fig. 3C).

### Mobility potential of antibiotic resistome in culture-enriched carbapenem and cephalosporin resistant microbiomes

To explore the spread potential of antibiotic resistance via horizontal gene transfer (HGT), ARGs-carrying contigs were constructed and identified from culture-enriched metagenomes (n = 2521, N50= 42482 bp) and wastewater and river water (n = 174, N50= 2895 bp) metagenomes, respectively (Table S5). A total of 2521 ARG-carrying resistance contigs were recovered from 92.2 Gbp culture-enriched metagenomes (n = 15), while constrastingly only 174 resistance contigs (of which 108 were assembled from influent metagenomes) were constructed from 63.5 Gbp wastewater (n = 4) and river water (n = 2) metagenomes. In general, most types of ARGs (excluding bacitracin, MLS and multidrug) in culture-enriched metagenomes, i.e., aminoglycoside (66.7%), beta-lactam (35.2%), chloramphenicol (74.6%), polymyxin (39.6%), quinolone (48.9%), sulfonamide (45.2%), tetracycline (54.8%), and trimethoprim (82.6%), were considerably located on plasmids (Fig. S4), demonstrating the widespread occurrence of ARGs on plasmids and their spread potential via HGT in carbapenem and cephalosporin resistant microbiomes. Furthermore, 266 resistance contigs from culture-enriched metagenomes were identified to carry various MGEs (Table S6 and Fig. S5), accounting for a mobility incidence of 10.6% which is comparable to those found in Swiss WWTPs ^16^.

Of the 3080 ORFs identified as ARGs in culture-enriched metagenomes, more than half ORFs (1780) were assigned to multidrug efflux genes, and 239 ORFs were annotated as beta-lactamase genes, including 98 carbapenemase genes and 52 ESBL genes (Table S7). However, only 29 beta-lactamase genes were recovered from wastewater and river water metagenomes. In culture-enriched metagenomes, 37 beta-lactamase genes, including 8 carbapenemase genes and 8 ESBL genes, were co-localized with diverse MGEs on 36 resistance contigs (Table S6), with a mobility incidence of 15.5%. For example, clinically-significant carbapenem resistance genes, such as KPC-2 (in contig KIN_1816 and CIN1_3707) and NDM-6 (in contig KIN_3253 and KSE100_4007), were identified to co-locate with transposase genes (Fig. 4A). Most strikingly, some carbapenem resistance genes (i.e, NDM-6 in contig CFE500_173) and polymyxin resistance genes (i.e., MCR-1 and MCR-4 in contig CSE100_19 and CUS100_5495, respectively), were presumably located on conjugative plasmids (Fig. 4B), as genes involved in plasmid conjugation (i.e., relaxase and type IV secretion system) were also identified in the same contigs (Table S6).

**Fig. 4.**
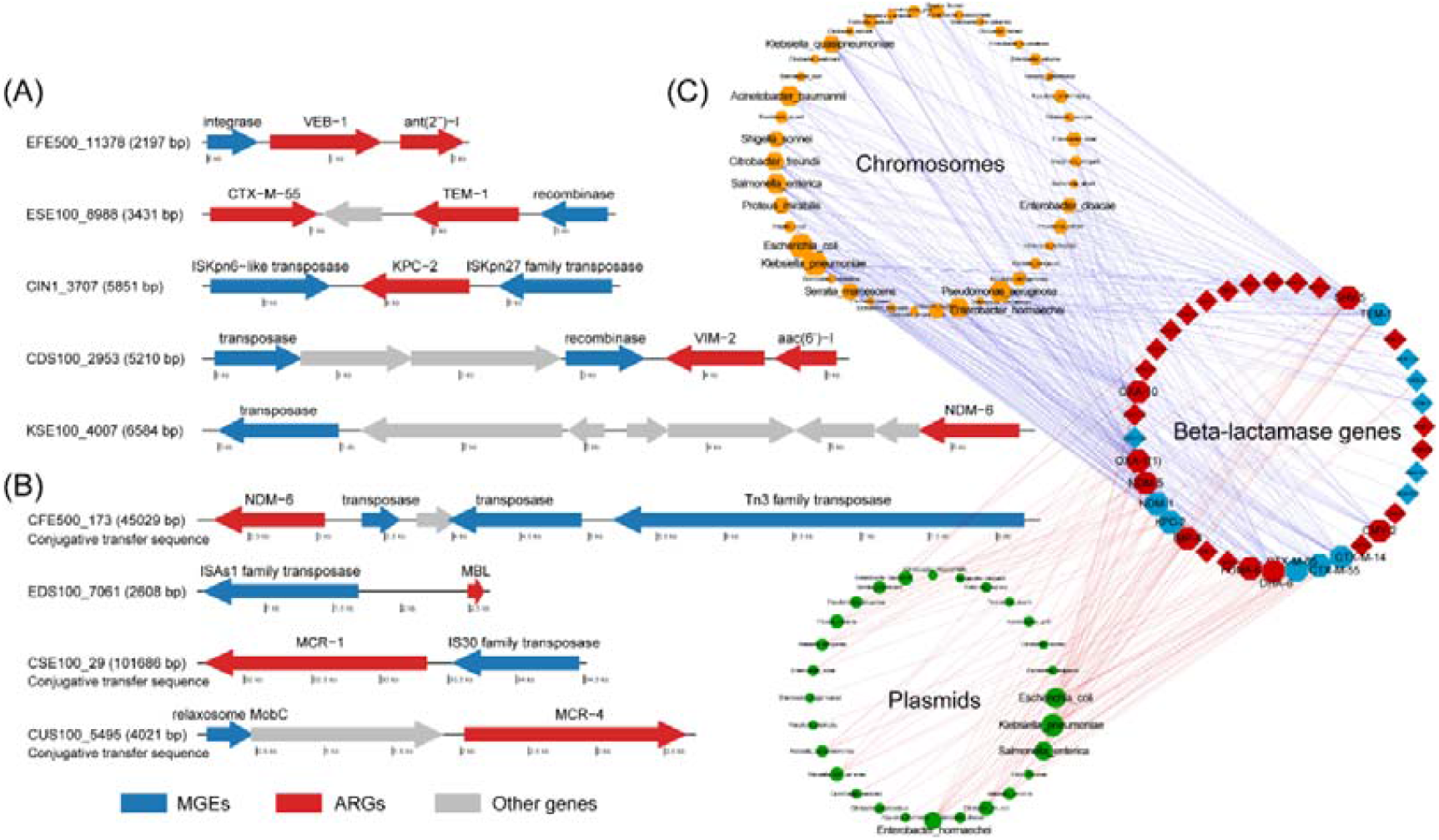
Mobility potential of representative ARGs from culture-enriched microbiomes and their relationship with gene counterparts in clinical-isolated bacterial pathogens. (A), Representative beta-lactamase genes flanking with MGEs, which share 100% nucleotide identity with genes in clinical-isolated bacterial pathogens. (B), Other representative beta-lactamase genes and polymycin resistance genes located on plasmid and/or flanking with MGEs. The contig ID is composed of the sample ID and contig serial number. The sample ID explanation is the same as that in the caption of Fig. 1. (C), Beta-lactamase genes from culture-enriched microbiota that shows 100% nuleotide sequence identity with genes in clinical-isolated bacterial pathogens. Octagons denote those beta-lactamase genes from culture-enriched microbiota which have abundant connections with clinical pathogens and blue octagons denote genes flanked by MGEs. Orange hexagons and green circles denote clinical pathogens’ chromosomes and plasmids, respectively. Node size represents the number of chromosomes or plasmids of HBP isolates, which was log_10_ normalized, and the edge connects beta-lactamase gene and the isolate harboring the identical gene counterpart.

We further explored the transfer potential of beta-lactamase genes between culture-enriched microbiomes and clinical HBPs, based on identical nucleotide sequence match ^19^. A total of 123 full-length and non-redundant (unique) beta-lactamase genes were blasted against the beta-lactamase database derived from 56882 genomes of clinical HBP isolates collected (NCBI Pathogen Detection). As a result, 38 beta-lactamase genes showed 100% global nucleotide sequence identity with a total of 6475 beta-lactamase genes on chromosomes or plasmids of 5084 clinical HBPs from 18 genera (Fig. 4C). Of the 38 beta-lactamase genes classified as carbapenemases (10), ESBLs (17), and other beta-lactamases (11), almost one third (12 genes) were flanked with MGEs (Table S8), which was 2-fold higher than the mobility incidence of the total 239 beta-lactamase genes (15.1%). *Escherichia coli* (1572 genomes, hereinafter), *Klebsiella pneumoniae* (2227), *Acinetobacter baumannii* (339), *Pseudomonas aeruginosa* (455), *Salmonella enterica* (320) and *Enterobacter hormaechei* (277) are the dominant species that share identical beta-lactamase genes in their chromosomes with culture-enriched microbiomes. *Escherichia coli* (136), *Klebsiella pneumoniae* (289) and *Salmonella enterica* (70) are also the main species that share identical plasmid-borne beta-lactamase genes with culture-enriched microbiomes. Note that, the aforementioned bacterial species mostly belong to the notorious ESKAPE panel and the World Health Organization priority pathogens list ^20,21^. Overall, the results showed complex beta-lactamase gene transfer network between or within clinical HBPs and culture-enriched microbiomes. For example, KPC-2, co-localized with two transposases, in contig CIN1_3707 had 100% similarity with chromosomal gene counterparts from 13 species (e.g., *Citrobacter freundii, Enterobacter hormaechei, Klebsiella pneumoniae, Pseudomonas aeruginosa, Salmonella enterica*) and plasmid-borne gene counterparts from 9 species (Fig. 4C). Moreover, 28, 13 and 12 of the 38 beta-lactamase genes were identified on chromosomes of different *Klebsiella pneumoniae, Pseudomonas aeruginosa* and *Salmonella enterica* genomes, respectively (Fig. 4C), indicating the great diversity of beta-lactamases in these known clinical HBPs and their wide interactions with culture-enriched microbiomes.

### Key features of ARG hosts including multi-antibiotic and polymyxin resistant *Enterobacteriaceae* predicted from phenotypic metagenome-assembled genomes

Besides the resistome composition and mobility, we further examined other key features related to health risks of ARGs including the genome-informed host taxonomy, pathogenicity, and multi-resistance. A total of 87 MAGs were recovered from culture-enriched samples with a quality over 50%, of which *Acinetobacter* (13), following by *Pseudomonas* (12), *Escherichia* (9), *Cupriavidus* (8), *Aeromonas* (7), *Klebsiella* (6), *Chryseobacterium* (5) were most dominant (Table S9). 55 unique MAGs were retained after dereplication and used for downstream analysis. These MAGs altogether accounted for 52.1% to 86.3% of paired metagenomic reads in culture-enriched metagenomes (Fig. S6A), representing most of the phenotypically selected bacterial communities.

We firstly checked the pathogenicity and resistome profiles of ARG hosts. Benefiting by the high resolution of genome-level taxonomy classification down to species level, we identified 19 of 55 MAGs recovered from culture-enriched microbiomes within the published reference pathogen lists ^22,23^, while 48 MAGs were annotated to carry a total of 3472 diverse virulence factors, such as invasion, adherence, and exotoxin (Table S10). However, only 102 virulence factors were identified in 38 of the 154 MAGs recovered from direct metagenomic sequencing, indicating that resistant pathogens were greatly enriched by selective agar media. Moreover, 43 MAGs were found carrying at least two ARGs, including 35 MAGs carrying multiple ARG types (Fig. 5A). The resistome profiles were distinct among bacterial species. In general, *Enterobacteriaceae* MAGs carried the most diverse ARGs types (7-11) and highest number of ARGs (29-49), following by some species of *Acinetobacter* MAGs, while *Pseudomonas* MAGs only harbored up to four types and ten numbers of ARGs (Fig. 5A). For example, an MAG annotated as *Escherichia dysenteriae* in culture-enriched microbiomes harbored 47 ARGs encoding aminoglycoside, bacitracin, beta-lactam, MLS, polymyxin and quinolone resistance and multidrug efflux pumps. Remarkably, polymyxin resistance genes were identified in all *Enterobacteriaceae* MAGs, i.e., *Escherichia* species, *Klebsiella pneumoniae, Enterobacter kobei* and *Citrobacter freundii*. Of greatest concern, *Enterobacter kobei* and *Citrobacter freundii* MAGs also harbor carbapenemase genes (i.e., NDM-6 or NDM-16) and other ARGs conferring resistance to aminoglycoside, MLS, quinolone, etc. The *Enterobacteriaceae* MAGs, including *Escherichia dysenteriae, Klebsiella pneumoniae, Enterobacter kobei* and *Citrobacter freundii*, were also characterized to share identical beta-lactamase genes with diverse species of clinical bacterial HBPs (Table S11). For example, SHV-5 ESBL gene of *Klebsiella pneumoniae* MAG was also identically encoded by chromosomes or plasmids of 22 HBPs species, such as *Salmonella enterica, Raoultella planticola, Proteus mirabilis, Acinetobacter baumannii* (Fig. 5B). *Citrobacter freundii* MAG surprisingly shared identical NDM-6 carbapenemase gene with chromosomes or plasmids of 16 HBPs species including species from distant phylogenetic lineages, e.g., *Pseudomonas aeruginosa, Acinetobacter baumannii* and *Providencia rettgeri*. In contrast to culture-enriched metagenome, only 18 ARGs carried by 13 MAGs were identified from 154 MAGs recovered from wastewater and river water metagenomes (Table S12). Thus, direct metagenomic sequencing could not enable a comprehensive or more complete profile of clinically important ARG hosts in environmental samples (e.g., wastewater and river water examined in this study), as ARG hosts are overwhelmed by huge genetic background of environmental microbiomes.

**Fig. 5.**
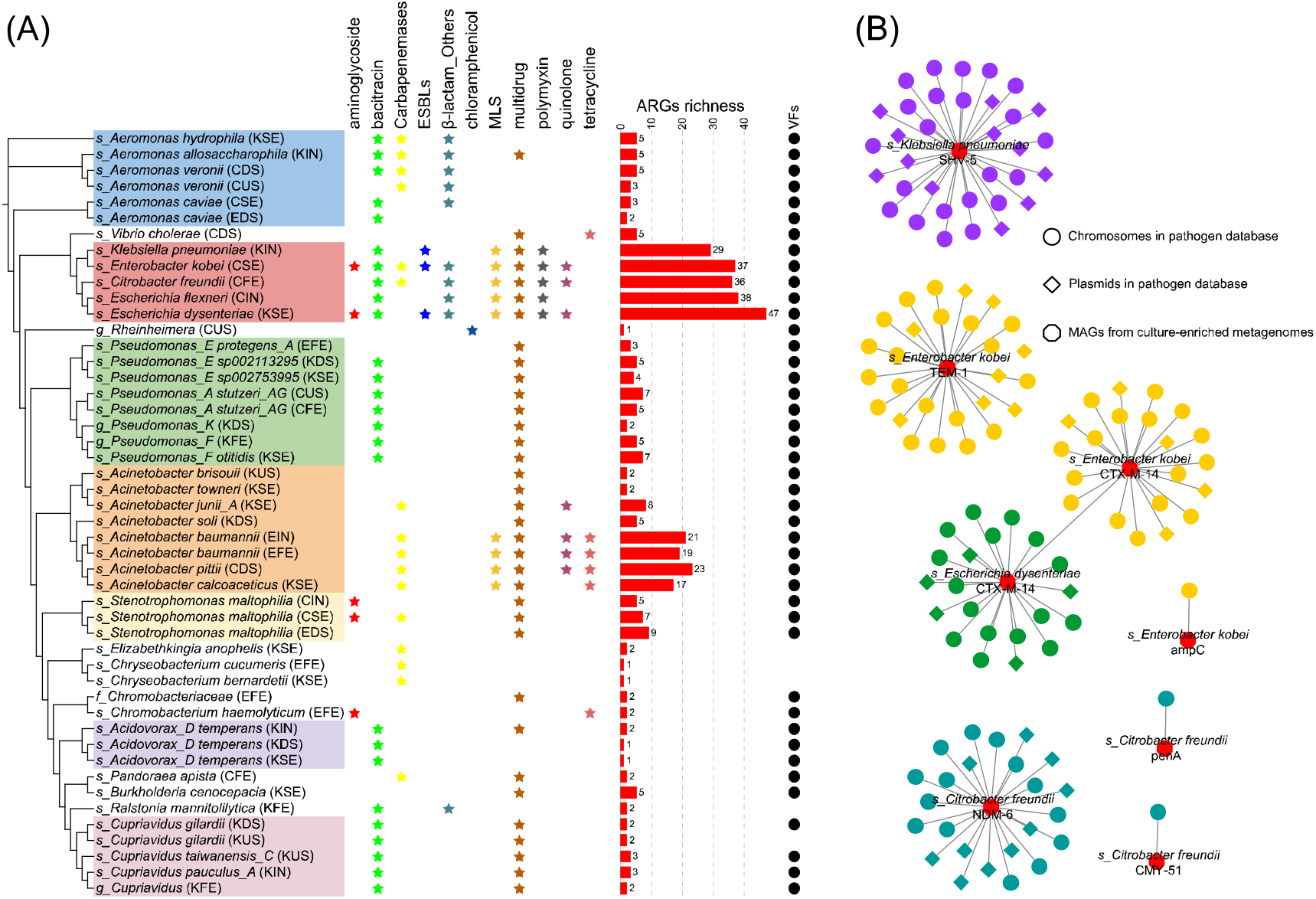
Key features of metagenome-assembled genomes (MAGs) recovered from culture-enriched microbiomes. (A), Phylogenetic distribution of 48 high-quality MAGs. The phylogenetic tree is constructed by FastTree using 120 bacterial marker genes from GTDB and visualized in iTOL. The first and latter two alphabets in the bracket denote agar types and sampling sites, respectively. Corlored asterisks denote ARG types carried by the corresponding MAG, the red bars display the total number of ARGs carried by the corresponding MAG, and the black circles indicate MAGs with virulence factors. The letter ‘s’,’g’,’f’ denotes species, genus and famly level, respecitvely, which represents the lowest assignable taxnomic rank of an MAG. (B), Networks of MAGs from culture-enriched metagenomes and clinical HBPs. Each node represents an MAG (octagon), chromosome (circle) or plasmid (rhombus) of the bacterial species. Edges connect species that share identical beta-lactamase genes.

### Mitigation of carbapenem and cephalosporin resistant bacteria co-selected with multi-antibiotic resistance

High-quality MAGs analysis enables the quantitative tracking of ARG hosts during wastewater treatments based on reads mapping to MAGs. The results showed that multi-antibiotic resistant *Enterabacteriaceae* and *Acinetobact*er in culture-enriched secondary effluent were still as abundant as those in culture-enriched influent, but greatly removed by the tertiary treatment (V-type sand filter), (Fig. S6A). Conversely, extended-spectrum beta-lactam resistant bacteria encoding less ARGs (e.g., *Pseudomonas, Cupriavidus*, or *Aeromonas*) superseding *Enterabacteriaceae* and *Acinetobact*er became the dominant ones. For example, the relative abundance of *Enterobacteriaceae* cultured on the KPC agar, CRE agar and ESBL agar greatly decreased from 0.88%, 22.6% and 72.6% in influent to 0.08%, 0.35% and 19.2% in downstream river, respectively, while *Pseudomonas* cultured on KPC agar, CRE agar and ESBL agar increased from 33.6%, 24.3% and 0.39% in influent to 44.4%. 56.55% and 35.8% in downstream river, respectively. Hence, the shift of bacterial community composition contributed to the decrease of overall ARGs level in culture-enriched microbiomes. For example, the reduction of MLS and polymyxin resistance genes in culture-enriched metagenomes, as shown in Fig. 3A, was attributed to the mitigation of their hosts, i.e., *Acinetobacter* or/and *Enterobacteriaceae*, by wastewater treatments (Fig. S6A), as inferred by the MAG-conferred linkages between ARGs and host taxonomy (Fig. 5).

Besides, by mapping metagenomic reads of wastewater and river water metagenomes to MAGs recovered from culture-enriched metagenomes, we revealed that carbapenem and cephalosporin resistant bacteria (e.g., *Enterobacteriaceae* and *Acinetobacter*) were remarkably mitigated by two orders of magnitudes in relative abundance by wastewater treatments (Fig. S6B). For instance, carbapenem and cephalosporin resistant *Enterobacteriaceae* were removed from 0.41% in influent to 0.02% in secondary effluent and further to 0.006% by wastewater disinfection treatment (equal to background abundance of 0.006% in receiving river water). Thus, both culture-enriched and direct metagenomic sequencing analyses all advocate that advanced treatment processes (e.g., tertiary filtration treatment and UV disinfection) of secondary wastewater should be enhanced in WWTPs, which is essential for the control of clinically relevant pathogens and antibiotic resistance, especially for municipal ones that continuously receive sewage carrying diverse multi-antibiotic resistant HBPs.

## Discussion

Municipal WWTPs and their receiving rivers, which are critical components of urban water cycles, continuously receive anthropogenic disturbances especially faecal pollution ^24^. They are under intensive scrutiny on the environmental occurrence and spread of clinically relevant antibiotic resistance ^25^, of which the surveillance of carbapenem and cephalosporin resistant bacteria is of the highest priority because of their high clinical importance ^26^. However, the high bacterial diversity and community complexity (e.g., WWTPs ^27,28^) makes it difficult to monitor clinically-important but usually rarely-present extended-spectrum beta-lactam resistant bacteria in environmental reservoirs. In this article, we proposed the use of culture-enriched phenotypic metagenomics to not only magnify signals of phenotypically selected microbes, but also largely simplify microbial diversity in inoculating wastewater and river water samples, as recently noticed when applied for human gut and lung samples ^29,30^. Based on 16S rRNA gene ASV analysis of culture-enriched agar microbiomes, bacterial composition in the original wastewater and river water samples was greatly simplified by phenotypic screening with extended-spectrum beta-lactams (i.e., on average 279 ASVs vs. 1804 ASVs before and after screening), and a targeted fraction of microbiomes was therefore greatly enriched, leading to the successful detection of clinically-important but usually rarely detectable carbapenem and cephalosporin resistant bacteria. For instance, we achieved high-throughput detection of 163 bacterial ASVs of *Enterobacteriaceae* accounting for up to 68.7% culture-enriched microbiomes inoculating from wastewater and river water samples, in which they mostly occurred as rare populations (a total abundance of 0.04% to 5.0%) that would have escaped detection without culture enrichment and phenotypic screening. At such a low abundance, direct metagenomic sequencing could not fetch enough genetic information to fulfil their qualitative and quantitative analyses, whereas phenotypic selection well overcame this outstanding technical bottleneck thanks to their prominent enrichment (by 13 to 1717 times) in the culture-enriched metagenomes. The presence of various clinically important carbapenem or cephalosporin resistant HBPs in culture-enriched samples was also validated by MALDI-TOF biotyping, discerning isolates down to species level.

Culture-enriched phenotypic metagenomics detected greater diversity and number of ARGs compared with direct metagenomic sequencing on both reads level and contig level. In total, 239 diversified beta-lactamase genes were assembled from culture-enriched metegenomes, which enables comprehensive insights into their mobility potential, clinical importance, and health risks. In contrast, it is difficult for direct metagenomic sequencing to achieve the above goals, as only limited number of beta-lactamase genes (i.e., 29 genes) were recovered from wastewater and river water metagenomes, even at much higher sequencing depths than culture-enriched metagenomes. Moreover, culture-enriched phenotypic metagenomics revealed some NDM family carbapenemases (e.g., NDM-6) identified in this study located on the conjugative plasmids (Fig. 4B). Their diversified variants have now been widely characterized ^5^ and found to associate with at least 20 different plasmid types ^31^. In this study, NDM family was also among the most dominant carbapenemase genes in the WWTP and its receiving river (Fig. 3C), which also comprises the most clinically significant carbapenemases ^4^. Despite that KPC family carbapenemases, as one of the most frequent carbapenemases in nosocomial environments ^4^, showed a relatively low abundance (up to 0.01 GPC) in culture-enriched microbiomes, two KPC family carbapenemases, flanking with transposases, putatively identified as plasmid sequences, were still recovered from culture-enriched metagenomes (Table S6). ARGs, especially those clinically-important ones, located on conjugative plasmids or flanked by MGEs, may cause greater environmental and health risks due to their strong HGT competence ^32^. CTX-M family was the most dominant ESBLs on culture-enriched microbiomes, followed by TEM, OXA and SHV family ESBLs (Fig. 3E). Clinical surveillance in many countries showed that CTX-M family ESBLs were replacing TEM and SHV family ESBLs as the predominant ESBLs in nosocomial isolates over the last decades ^33,34^. The consistent pattern strongly implicates an intimate resistome connection between nosocomial environments and WWTPs.

The evidence for ARGs communication between nosocomial HBPs and WWTP microbiomes was indeed identified in this study based on comparative nucleotide identity analysis, that is, 38 of 123 full-length beta-lactamase genes, including KPC, NDM, CTX-M, etc., from culture-enriched microbiomes were detected exactly the same as those encoded on the chromosome or/and plasmids of clinical HBP isolates assigned broadly to 18 bacterial genera (Fig. 4C). The hosts of the 38 beta-lactamase genes in the WWTP might directly from nosocomial environments, or ARGs transfer events between WWTP microbiome and clinical HBPs have recently happened, regardless of whether ARGs were moving from WWTPs to the clinic, or vice versa. Considering the strong mobility incidence of the 38 beta-lactamase genes (31.58%), which was 2-fold higher than the average level of beta-lactamase genes revealed in this study (Table S6), their presence, especially carbapenemase and ESBL genes, in the WWTP underscores the clinical importance of WWTP resistome, and emphasizes the alarming health risk with long-standing wastewater dissemination of clinically significant ARGs.

Culture-enriched phenotypic metagenomics also makes the high-quality genome reconstruction successful for culturable but relatively less abundant microbial populations with targeted resistance phenotypes (e.g., *Enterobacteriaceae* and *Acinetobacter*, less than 1.5% in influent microbiomes), and enables further comprehensive insights into the taxonomic identity, pathogenicity, and resistome profiles of ARG hosts on genome level for comprehensive assessment on their health risks. For instance, genome-level analysis revealed that *Enterabacteriaceae* MAGs harbored dozens of ARGs encoding resistance to over 10 classes of antibiotics, suggesting the alarming prevalence of co-resistance to numerous other antibiotics in the cultured carbapenem and cephalosporin resistant *Enterabacteriaceae* (Fig. 5). Note that, multi-antibiotic resistance should considerably facilitate the survival and dissemination of their bacterial hosts in environmental systems with anthropogenic stressors (e.g., antibiotics and heavy metals), such as WWTPs ^35^. High-quality MAGs, bridging taxonomy information and ARG profiles, also realizes the characterization of potential HGT events, as revealed in this study that distant phylogenetic lineages shared identical beta-lactamase genes. Most strikingly, carbapenemase genes and polymyxin resistance genes, locating on different contigs, were identified to simultaneously encoded by at least two MAGs, taxonomically assigned to *Enterobacter kobei* (MCR-1 and NDM-6) and *Citrobacter freundii* (ArnA and NDM-16), which would otherwise be overlooked if based on solely contig-level analysis. The co-presence of carbapenem and polymyxin resistance HBPs would take patients back to a “pre-antibiotic era”, as polymyxins, including colistin, are the last-resort treatment for serious nosocomial infections caused by carbapenem resistant bacteria ^36^. Further, our identification of carbapenem and polymyxin resistant HBPs in the single WWTP might launch alarm on their prevalence in other global regions and inspire efforts to examine its potential link with the increasing co-resistance to carbapenem and polymyxin in clinical *Enterobacteriaceae* isolates ^37,38^.

Despite the key importance of bridging the genotypes and phenotype gaps, it should be noted that resistance phenotypes observed may not always consistent with reported genotypes. For example, while *Escherichia* species and *Klebsiella pneumoniae* are widely reported typical carbapenemase producers, their recovered genomes did not contain any known carbapenemase genes. Instead, other beta-lactamase genes known as ESBLs or narrow-spectrum beta-lactam resistance genes (e.g., AmpC), plus numerous genes encoding various families (especially MFS and RND) of multidrug efflux pumps were identified, which could contribute to carbapenem resistance as previously reported ^3^. Therefore, carbapenem-resistant phenotype is not necessarily ascribed explicitly to known genotypes, and future functional genomics attempts to identify unreported or new resistance genes ^39,40^ or transcriptomic studies to check on activities of alternative resistance mechanisms ^16^ are warranted. In addition, MAG-based analyses tend to better represent chromosomal ARGs while underestimated plasmid ARGs. This might be one of the causes for the observed inconsistence between examined phenotypes and genotypes of some MAGs, as contig-based analyses supported the location of the relevant ARGs on plasmids.

Collectively, our study demonstrated the following technical advantages of culture-enriched phenotypic metagenomics. First, culture-enriched phenotypic metagenomics could realize fast and efficient profiles of the resistomes of microbiomes with specific resistance phenotypes. The heavy work burdens from conventional PCR of clinical isolates which is low-throughput, labor-intensive and high-cost was spared and replaced by culture-enriched metagenomic sequencing and coupled bioinformatic analysis as demonstrated here. Second, culture-enriched phenotypic metagenomics can enable us to simultaneously decipher the targeted living fraction of microbiomes with a given screening phenotype and their global genetic characteristics and functional potentials based on recovered high-quality MAGs with accurate taxonomic resolution of targeted species. In general, high-quality genomes are essential for bridging reliable linkage between antibiotic-resistance genotypes, phenotypes, and host pathogenicity and taxonomy, so as to get precise and credible risk ranking in environmental (e.g., wastewater) resistome with regard to better-informed infectious disease control in humans ^25,41^. Last, phenotypic metagenome-assembled genomes (MAGs) can be quantified by mapping the reads from direct metagenomic sequencing to reveal the fate of phenotypically screened and positively selected microbial lineages in environmental samples, habitats, and ecosystems. Promisingly, phenotypic metagenomics complemented with microfluidics, single-cell tagging, and long-read sequencing would realize high-throughput phenotypic screening and efficient genome reconstruction in a cheaper, faster, and convenient way and make the culture-enriched phenotypic metagenomics framework proposed here more powerful for antibiotic resistance and resistome surveillance.

## Methods

### Sample collection and pretreatment

The sampling campaign was conducted in a municipal WWTP with tertiary treatment and its receiving river located in Hangzhou, China. Six samples were collected in May 2020, including the influent (IN), secondary effluent (SE), V-type filter effluent (FE), disinfection effluent (EF), upstream (US) and downstream (DS) river water (Fig. 1). For each sampling site, 10-L water sample was taken and transported to the laboratory for further processing within 2 hours. Microbial solids were collected by filtering each water sample through 0.22-μm cellulose nitrate membranes. The obtained biomass-carrying membranes were further applied for phenotypic screening or stored at −20 °C before direct DNA extraction. The volumes of filtered wastewater and river water samples were listed in Table S1.

### Phenotypic screening and MALDI-TOF biotyping

For phenotypic screening and culture enrichment of targeted fraction of environmental microbiomes, the biomass-carrying membranes were incubated in triplicate for each sample on three different selective agars (CHROMagar™, France), that is, the KPC agar for detection of Gram-negative bacteria with a reduced susceptibility to most of carbapenem antibiotics, mSuperCARBA (CRE) agar for detection of Carbapenem-resistant Enterobacteriaceae, and ESBL agar for detection of gram-negative bacteria producing extended-spectrum beta-lactamases, which are widely applied for phenotypic screening of clinical carbapenem and cephalosporin resistance. The membrane-inoculated agars were cultivated at 35°C for 24 hours, and then the culture-enriched microbiomes carried by membranes were harvested for subsequent applications.

Matrix-assisted laser desorption/ionization time-of-flight mass spectrometry (MALDI-TOF MS) was used for bacterial taxonomy identification. Bacterial colonies were randomly picked from the above obtained membranes and re-cultured to confirm their resistance capacity by streak plate method. The purified colony materials were smeared onto a 384-well steel target plate using 10-µL pipette tips and then overlaid with 1 μL of HCCA matrix solution. *Escherichia coli* DH5α peptide with a protein profile was used to calibrate the instrument. Mass spectra of each isolate was automatically collected and analyzed by Microflex LT MALDI-TOF MS unit and MALDI BioTyper software, version 4.0 (Bruker Daltonics, Germany). The identification is based on comparisons between the mass spectra of the analyzed sample and the database.

### Microbial biomass collection and DNA extraction

Culture-enriched microbiomes were collected from agar plates by pooling triplicate filter membranes. Excluding cultured membranes of final effluent (EF) samples which carry no observable colony with 500-mL EF wastewater inoculated (Table S2), 30 agar cultures originating from 5 wastewater and river water samples with 3 agar media and 2 water filtering volumes were obtained. Detailed information was shown in Table S1. A total of 36 samples including 6 raw samples before cultivation and 30 cultured samples were applied for DNA extraction. Genomic DNA was extracted using FastDNA Spin Kit for Soil (MP Biomedicals, USA) following the manufacturer’s instructions, and was applied for 16S rRNA gene amplicon sequencing and metagenomics sequencing (Table S1).

### 16S rRNA gene amplicon sequencing and data processing

To explore microbial community composition of raw samples and culture-enriched samples, the V3-V4 hypervariable regions of bacterial 16S rRNA genes ^42^ were amplified using the forward primer 341F (5’-CCTACGGGNGGCWGCAG-3’) and reverse primer 802R (5’-GACTACHVGGGTATCTAATCC-3’), and then the amplicons were sequenced on the Illumina Miseq platform (PE250) of the Novogene Corporation (Beijing, China). The 16S rRNA gene amplicon sequencing data was processed using Qiime2 pipeline (v2020.6.0) ^43^ and DADA2 algorithm (v2020.6.0) ^44^. After quality trimming, merging of paired sequences and removal of chimeric sequences, an abundance table of amplicon sequence variant (ASV) was generated based on 100% sequence similarity for 16S rRNA genes. Taxonomic assignment of the ASVs was conducted using the Silva database (version 132) ^45^. The final ASV table containing ASV id, taxonomic classification and abundance information was used for following analysis.

### Metagenomics sequencing and data processing

The genomic DNA was sent to the Novogene Corporation (Beijing, China) for metagenomics sequencing on the Illumina’s NovaSeq 6000 platform with a paired-end 150 bp strategy. On average, 21.7 million and 36.6 million raw reads were generated for each culture-enriched sample and raw sample, respectively. The detailed information of metagenomic datasets was shown in Table S3.

For each metagenomic data, raw reads were filtered to remove low-quality reads and adapters using Trimmomatic (v0.39) ^46^. Clean reads were then de novo assembled using metaspades option in MetaWRAP pipeline (v1.3.0) ^47^ with default parameters. Assembly results were summarized in Table S3. Open reading frames (ORFs) were predicted for each contig using Prodigal (v2.6.3) ^48^. The ARG-like ORFs were identified by aligning their nucleotide sequences to the SARG database (v 2.0) ^18^ using BLASTX at E value ≤ 10^−7^ with at least 80% similarity and 70% query coverage. The ORFs located on ARG-carrying resistance contigs were then annotated against the bacterial sub-library of NCBI non-redundant (NR) protein database (retrieved on July 26, 2020) using BLASTP at E value ≤ 10^−5^ with the above threshold. MGEs carried by resistance contigs were searched based on string matches of the NR annotation results, as described previously ^49^. Co-occurrence of ARGs and MGEs was considered positive if an ARG was found within ten open reading frames from upstream or downstream a MGE on the same contig ^50^. Further, the mobility potential of a ARGs cassette was quantified by mobility incidence (M%) as proposed by Ju et al. ^16^. The location (plasmid or chromosome) of resistance contigs were predicted by PlasFlow (v1.1) ^51^ with default parameters.

Contigs from each sample were binned independently to generate metagenome-assembled genomes (MAGs) using three different binning software (i.e., MetaBAT2, MaxBin and CONCOCT) in MetaWRAP pipeline (v1.3.0) ^47^ with default parameters. Then high-quality draft genomes were extracted from the above generated MAGs using bin refinement module, and further bin reassembly module was applied to improve the MAGs’ quality by enhancing the completeness and reducing the contamination. Only MAGs with an overall quality ≥50% (completeness – 5×contamination) were retained for downstream analysis. Finally, a total of 87 high-quality MAGs were obtained from culture-enriched microbiome. MAGs were taxonomically assigned using classify_wf module of gtdbtk (v1.4.0), phylogenetic analysis of MAGs was conducted with gtdbtk infer module based on a set of 120 bacterial specific marker genes from GTDB ^52^, and the phylogenetic tree was visualized in iTOL ^53^. Virulence factors in MAGs were identified by against the core database of VFDB (retrieved on Jan. 14, 2022) ^54^ using BLASTX at E value ≤ 10^−7^ with at least 80% similarity and 70% query coverage.

To calculated the relative abundance, the MAGs were firstly dereplicated using dRep (v3.0.0) with the ANI ≥ 99% in secondary cluster ^55^. Then these MAGs were merged to build an index file and the clean reads from each sample were mapped to the index file using Bowtie2 (v2.3.4.1) ^56^. Then, an in-house perl script (https://github.com/RichieJu520/Metagenomics-workshop-on-euler) was used to calculate the relative abundance of MAGs. Briefly, the relative abundance of each MAG was calculated as the number of reads aligning to the MAG normalized by the total number of reads in the sample.

## Data Availability

The sequencing raw data, including 16S rRNA gene amplicon sequencing and metagenomic sequencing data, are accessible in the China National GeneBank (CNGB) with accession number CNP0002692, and the data are released to NCBI and ENA from CNGB (accession numbers will be later provided upon allocation).

## Acknowledgments

This work was supported by Zhejiang Provincial Natural Science Foundation of China under Grant No. LR22D010001 and National Natural Science Foundation of China under Grant No. 51908467. We would like to thank Dr. Helmut Bergmann at EAWAG, Switzerland for methodology advice and helpful discussion.

