## Supplementary Information for "Using Culture-Enriched Phenotypic Metagenomics for Targeted High-Throughput Monitoring of Clinically-Important Fraction of Beta-Lactam Resistome"

**1. Supplementary tables**

**Table S1.** List of samples for 16S-V3V4 hypervariable region amplicon sequencing and metagenomic sequencing.

**Table S2.** KPC agar-, CRE agar- and ESBL agar-cultured bacterial concentration in the wastewater treatment plant and river water.

**Table S3**. The information of the metagenomic datasets and quality statistics of metagenomic assembly.

**Table S4.** Dominant types of beta-lactamase genes in clinical pathogens.

**Table S5.** Location of ARGs-carrying resistance contigs

**Table S6.** Co-occurrence of ARGs and MGEs in culture-enriched metagenomes

**Table S7.** Location of ARGs in culture-enriched metagenomes grouped by agar type

**Table S8.** Beta-lactamase genes sharing 100% sequence identity with gene countparts in clinical pathogens

**Table S9.** Summary of the ARG-carrying MAGs recovered from culture-enriched metagenomes

**Table S10.** Virulence factors in MAGs recovered from culture-enriched metagenomes and wastewater and river water metagenomes

**Table S11.** Summary of MAGs that share identical bate-lactamase genes with clinical HBPs.

**Table S12.** ARGs in MAGs recovered from wastewater and river water metagenomes

Table S3, S5, S6, S7, S8, S9, S10, S11 and S12 in the Supplementary Datasets.

**2. Supplementary figures**

**Fig. S1.** Bacterial community composition of the culture-enriched *Enterobacteriaceae* on the KPC, CRE and ESBL agar.

**Fig. S2.** Observed ARG subtypes in culture-enriched samples and original water samples.

**Fig. S3.** Representative co-occurrence of ARGs revealed by gene arrangements.

**Fig. S4.** The genetic locations of ARG-carrying contigs.

**Fig. S5.** Number of different types of MGEs detected across all culture-enriched metagenomes.

**Fig. S6.** Comparison of the total relative abundance of dereplicated MAGs among (A) culture-enriched samples and (B) wastewater and river water samples.

**Table S1.** List of samples for 16S-V3V4 hypervariable region amplicon sequencing and metagenomic sequencing.

| Sample | 16S V3V4 | Metagenomic | Volume (mL) |  | Sample | 16s V3V4 | Metagenomic | Volume (mL) |
| --- | --- | --- | --- | --- | --- | --- | --- | --- |
| KIN1 | ● | ● | 1 |  | EIN1 | ● | ● | 1 |
| KIN0.1 | ● |  | 0.1 |  | EIN0.1 | ● |  | 0.1 |
| KSE100 | ● | ● | 100 |  | ESE100 | ● | ● | 100 |
| KSE10 | ● |  | 10 |  | ESE10 | ● |  | 10 |
| KFE500 | ● | ● | 500 |  | EFE500 | ● | ● | 500 |
| KFE100 | ● |  | 100 |  | EFE100 | ● |  | 100 |
| KEF500 |  |  | 500 |  | EEF500 |  |  | 500 |
| KEF100 |  |  | 100 |  | EEF100 |  |  | 100 |
| KUS100 | ● | ● | 100 |  | EUS100 | ● | ● | 100 |
| KUS10 | ● |  | 10 |  | EUS10 | ● |  | 10 |
| KDS100 | ● | ● | 100 |  | EDS100 | ● | ● | 100 |
| KDS10 | ● |  | 10 |  | EDS10 | ● |  | 10 |
| CIN1 | ● | ● | 1 |  | IN | ● | ● | 60 |
| CIN0.1 | ● |  | 0.1 |  | SE | ● | ● | 1100 |
| CSE100 | ● | ● | 100 |  | FE | ● | ● | 3000 |
| CSE10 | ● |  | 10 |  | EF | ● | ● | 2500 |
| CFE500 | ● | ● | 500 |  | US | ● | ● | 350 |
| CFE100 | ● |  | 100 |  | DS | ● | ● | 350 |
| CEF500 |  |  | 500 |  |  |  |  |  |
| CEF100 |  |  | 100 |  |  |  |  |  |
| CUS100 | ● | ● | 100 |  |  |  |  |  |
| CUS10 | ● |  | 10 |  |  |  |  |  |
| CDS100 | ● | ● | 100 |  |  |  |  |  |
| CDS10 | ● |  | 10 |  |  |  |  |  |

**Table S2.** KPC agar-, CRE agar- and ESBL agar-cultured bacterial concentration (CFU/mL) in the wastewater treatment plant and river water.

|  | IN | SE | FE | EF | US | DS |
| --- | --- | --- | --- | --- | --- | --- |
| KPC agar | 1033.3 | 3.4 | 0.9 | 0 | >50 | >50 |
| CRE agar | 1046.7 | 1.4 | 0.4 | 0 | >50 | >50 |
| ESBL agar | >3000 | 7.1 | 0.7 | 0 | >50 | >50 |

Note: No colony formed on the three selective agars with 500-mL final effluent wastewater inoculated.

**Table S4.** Dominant types of beta-lactamase genes in clinical pathogens.

| Gene type | ORFs count | Ratio |
| --- | --- | --- |
| **OXA** | **20474** | **77504**/83834 |
| class C | 14812 |  |
| **TEM** | **14513** |  |
| **SHV** | **9611** |  |
| **CTX-M** | **8630** |  |
| **KPC** | **3828** |  |
| PDC | 2414 |  |
| **NDM** | **1909** |  |
| CMY | 1313 |  |

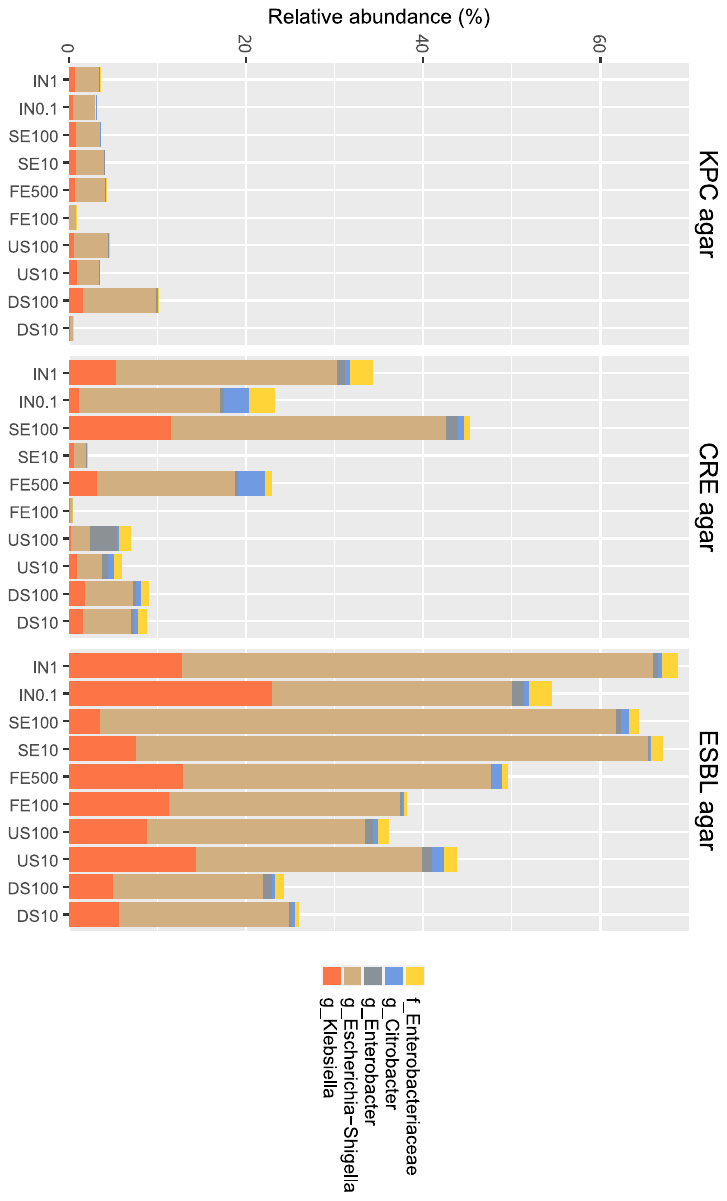

**Fig. S1.** Bacterial community composition of the culture-enriched *Enterobacteriaceae* on the KPC, CRE and ESBL agar.

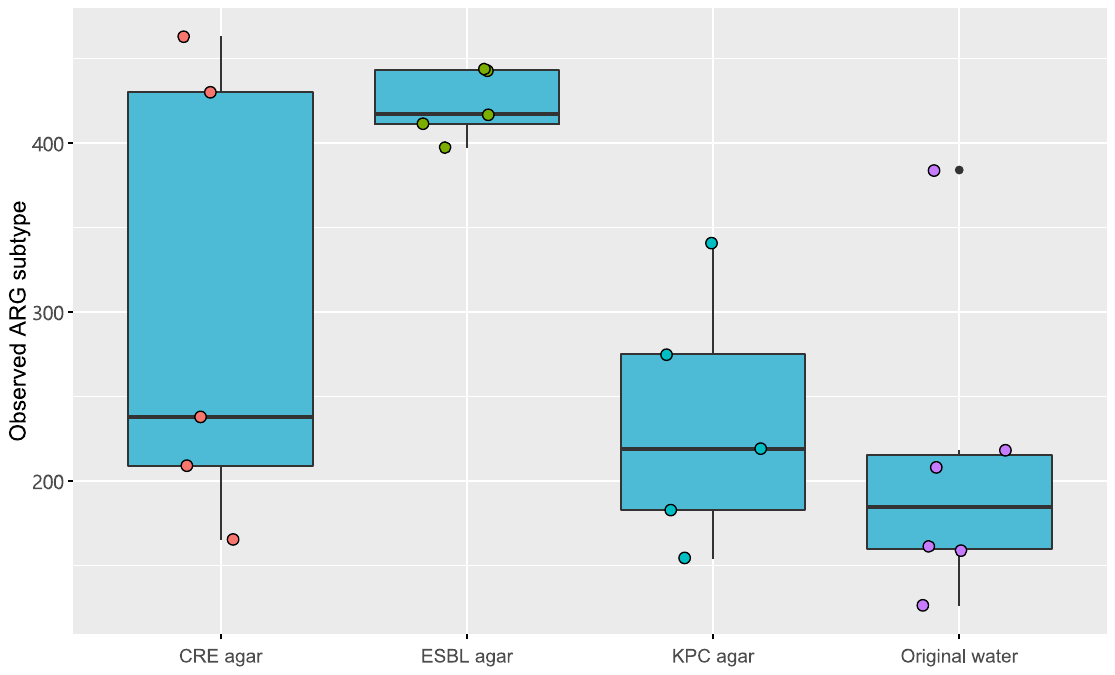

**Fig. S2.** Observed ARG subtypes in culture-enriched samples and original water samples. Original samples mean wastewater or river water samples without cultivation.

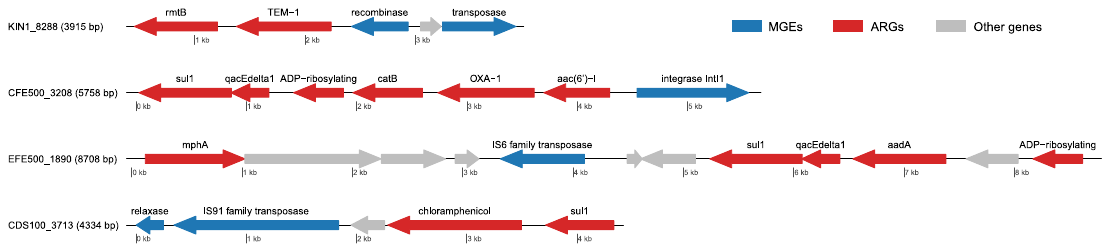

**Fig. S3.** Representative co-occurrence of ARGs revealed by gene arrangements.

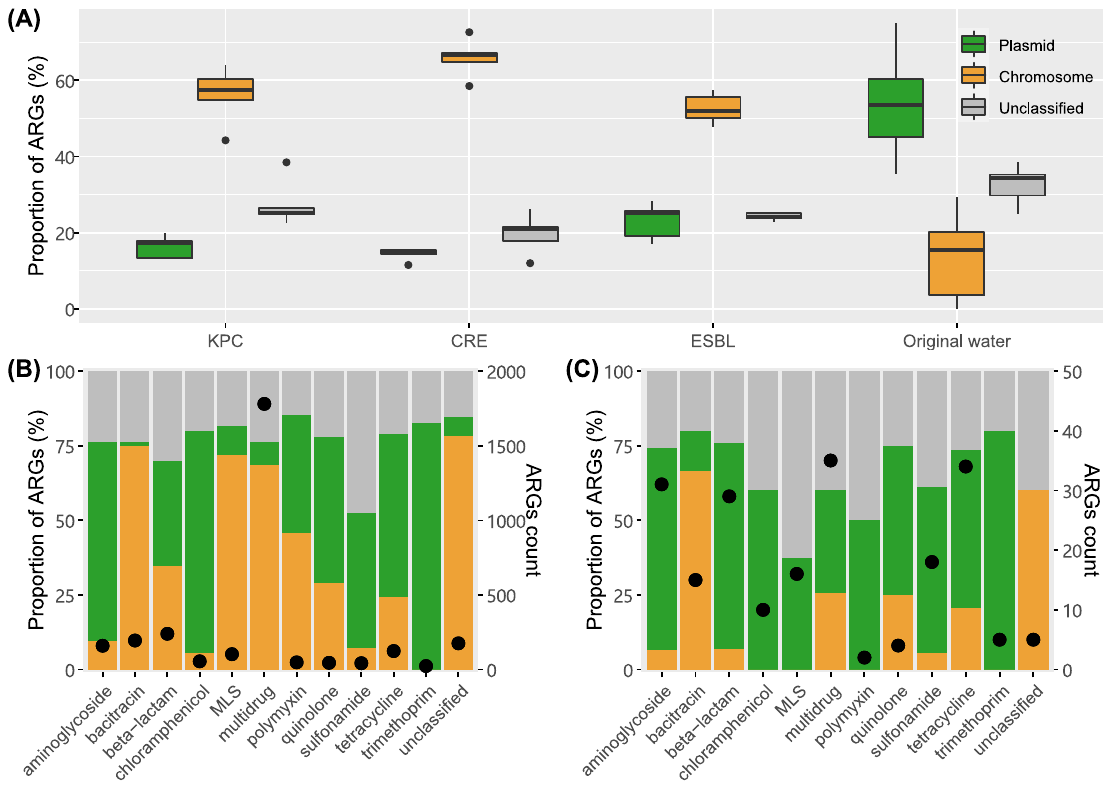

**Fig. S4.** The genetic locations of ARG-carrying contigs. (A), Comparison of the proportions of ARG-carrying contigs located on plasmids and chromosomes and unclassified sequences in culture-enriched micriobiota and original samples. Original water samples represent wastewater or river water samples without cultivation. (B) and (C), The proportions of different ARG types located on plasmid, chromosomal and unclassified sequences in culture-enriched metagenomes and wasterwater and river water metagenomes, respectively. MLS: macrolide-lincosamide-streptogramin. Green, orange and grey represent plasmid, chromosomal and unclassified sequences, respectively. MLS: macrolide-lincosamide-streptogramin.

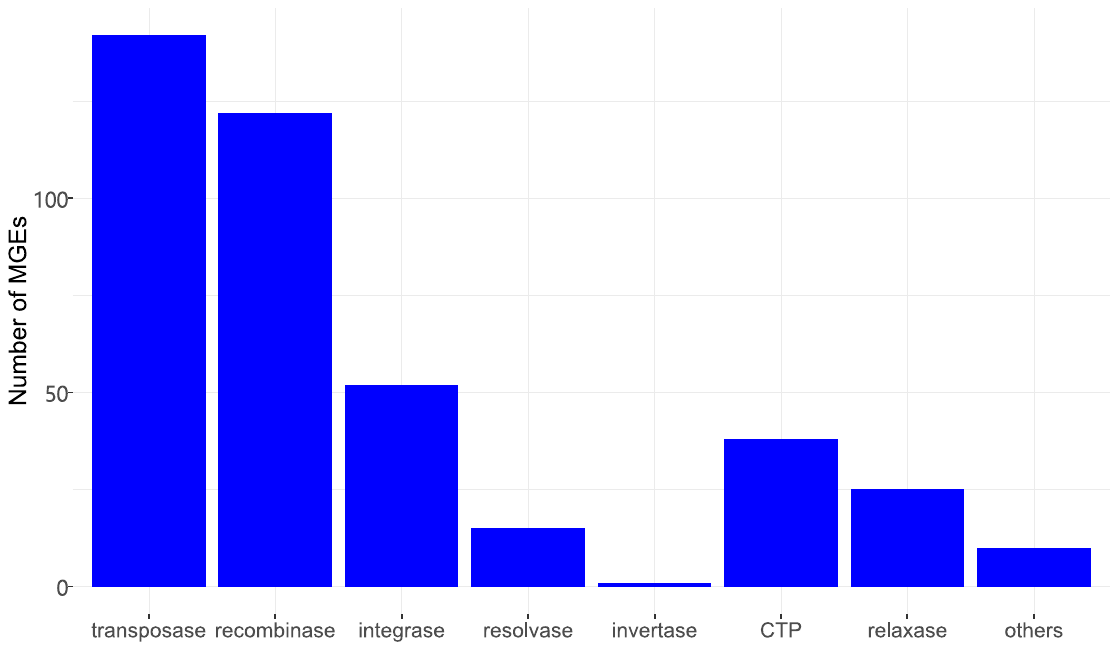

**Fig. S5.** Number of different types of MGEs detected across all culture-enriched metagenomes. CTP: Conjugal transfer protein.

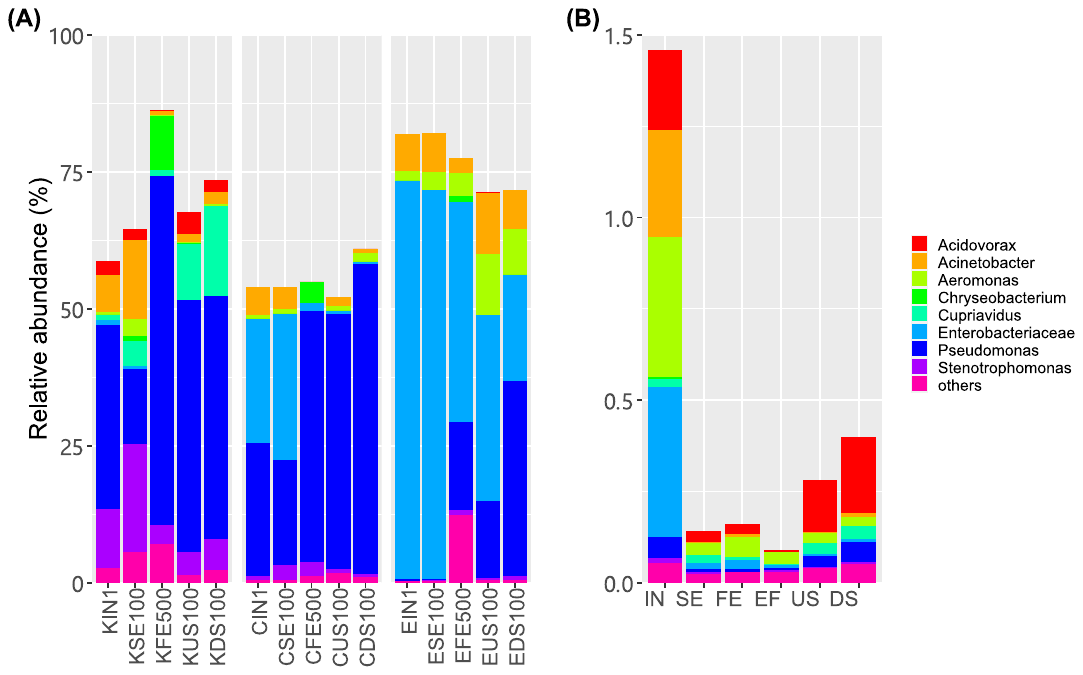

**Fig. S6.** Comparison of the total relative abundance of dereplicated MAGs among (A) culture-enriched samples and (B) wastewater and river water samples.
